# Prophylactic and Therapeutic Inhibition of In Vitro SARS-CoV-2 Replication by Oleandrin

**DOI:** 10.1101/2020.07.15.203489

**Authors:** Kenneth S. Plante, Jessica A. Plante, Diana Fernandez, Divya Mirchandani, Nathen Bopp, Patricia V. Aguilar, K. Jagannadha Sastry, Robert A. Newman, Scott C. Weaver

**Author notes:** Corresponding authors: Scott C. Weaver,; Kenneth S. Plante.

## Abstract

With continued expansion of the COVID-19 pandemic, antiviral drugs are desperately needed to treat patients at high risk of life-threatening disease and even to limit spread if administered early during infection. Typically, the fastest route to identifying and licensing a safe and effective antiviral drug is to test those already shown safe in early clinical trials for other infections or diseases. Here, we tested *in vitro* oleandrin, derived from the *Nerium oleander* plant and shown previously to have inhibitory activity against several viruses. Using Vero cells, we found that prophylactic oleandrin administration at concentrations down to 0.05 μg/ml exhibited potent antiviral activity against SARS-CoV-2, with an 800-fold reduction in virus production, and a 0.1 μg/ml dose resulted in a greater than 3,000-fold reduction in infectious virus production. The EC_50_ values were 11.98ng/ml when virus output was measured at 24 hours post-infection, and 7.07ng/ml measured at 48 hours post-infection. Therapeutic (post-infection) treatment up to 24 hours after infection of Vero cells also reduced viral titers, with the 0.1 μg/ml dose causing greater than 100-fold reductions as measured at 48 hours, and the 0.05 μg/ml dose resulting in a 78-fold reduction. The potent prophylactic and therapeutic antiviral activities demonstrated here strongly support the further development of oleandrin to reduce the severity of COVID-19 and potentially also to reduce spread by persons diagnosed early after infection.

**IMPORTANCE:** COVID-19, a pandemic disease caused by infection with SARS-CoV-2, has swept around the world to cause millions of infections and hundreds-of-thousands of deaths due to the lack of vaccines and effective therapeutics. We tested oleandrin, derived from the *Nerium oleander* plant and shown previously to reduce the replication of several viruses, against SARS-CoV-2 infection of Vero cells. When administered both before and after virus infection, nanogram doses of oleandrin significantly inhibited replication by up to 3,000-fold, indicating the potential to prevent disease and virus spread in persons recently exposed to SARS-CoV-2, as well as to prevent severe disease in persons at high risk. These results indicate that oleandrin should be tested in animal models and in humans exposed to infection to determine its medical usefulness in controlling the pandemic.

## INTRODUCTION

After its emergence in late 2019, an epidemic of coronavirus disease termed COVID-19 was first recognized in the city of Wuhan, China, then quickly spread to pandemic proportions (1). As of July 13, 2020, over 13 million cases had been identified with nearly every country involved worldwide, and over 570,000 fatalities. Hospitals and other health care systems were overwhelmed in Wuhan, Italy, Spain and New York City before cases peaked in these locations. The underlying virulence of this newly emerged coronavirus is particularly severe for older age groups and those with chronic pulmonary and cardiac conditions, diabetes, and other comorbidities. Other factors fueling morbidity and mortality have included limited supplies in personal protective devices and ventilators required for endotracheal intubation (2). With no licensed vaccines available, and antiviral drugs other than Remdesivir (3) showing limited or no efficacy in clinical trials (4–7), the only methods available to limit infection and disease are public health measures designed to limit human contact and transmission.

Typically, the fastest route to identifying and licensing a safe and effective antiviral drug is to test those already shown safe in early clinical trials for other infections or diseases. A few FDA-approved drugs that target human proteins predicted to interact with SARS-CoV-2 proteins show antiviral activity in vitro, and Lopinavir-ritonavir, hydroxychloroquine sulfate, and emtricitabine-tenofovir marginally reduced clinical outcomes of infected ferrets but did not significantly reduce viral loads (8). Several drugs including ribavirin, interferon, lopinavir-ritonavir, and corticosteroids, have been used in patients infected with the closely related *Betacoronaviruses* SARS (from 2002-2003) or MERS (emerging since 2012), with varying efficacy results (9).

The class of compounds known as cardiac glycosides consists of well characterized molecules such as digoxin, digitoxin, ouabain and oleandrin. The well accepted mechanism of action for these compounds lies in their ability to inhibit functioning of Na,K-ATPase which, in turn, alters ion flux across membranes (10). This explains, for example, the ability of this class of compounds to improve functioning of heart muscle in patients with congestive heart failure. In the last several decades additional therapeutic uses of compounds such as oleandrin have been recognized. These include the use of this unique molecule, and extracts of the oleander (*Nerium oleander*) plant, as effective therapeutics that range from cosmetic treatment of skincare problems (11) to the treatment of cancer (12–14). Oleandrin has been recognized as the active principle ingredient in oleander extracts used in Phase I and Phase II clinical trials of patients with cancer (15). These trials defined the pharmacokinetics of oleandrin and demonstrated that extracts containing this molecule could be given safely as an oral drug to patients without significant adverse events.

Less appreciated is the strong antiviral activity of this class of compounds. A recent review described antiviral activities of cardiac glycosides against cytomegalovirus, Herpes simplex virus, adenovirus, chikungunya virus, coronaviruses, respiratory syncytial virus, Ebola virus, influenza virus and human immunodeficiency virus (16). We recently reported that oleandrin, a unique lipid-soluble cardiac glycoside obtained solely from *Nerium oleander*, has strong antiviral activity against HIV-1 and HTLV-1 (17, 18). Importantly, in studies with HIV-1, a unique antiviral activity of oleandrin was observed in terms of producing viral progeny with a greatly diminished potential to infect new target cells (17). This decreased infectivity was due in part to a reduction in the envelope glycoprotein of progeny virus, which is essential for viral infectivity. We also reported that oleandrin treatment of cells infected with HTLV-1 resulted in reduced infectivity of progeny virus particles. Oleandrin inhibited virological synapse formation and consequent transmission of HTLV-1 *in vitro* (18). Collectively this research strongly suggested that oleandrin and defined plant extracts with this molecule have a strong potential against enveloped viruses. Because SARS-CoV-2, the cause of COVID-19, is also an ‘enveloped’ virus we reasoned that oleandrin could be effective in reducing viral load in infected individuals via production of progeny virus with a significantly reduced potential to infect new target/host cells.

Considering the emergency need to identify potential antiviral therapies for COVID-19, we tested oleandrin in SARS-C0V-2-infected Vero 81 (African Green monkey kidney) cells at doses ranging from 1 μg/ml down to 0.005 μg/ml. Here, we show the strong inhibitory profile of oleandrin in greatly reducing infectious virus production.

## Results

### Toxicity of Oleandrin

A range of oleandrin concentrations (1 μg/ml to 0.005 μg/ml) was selected on the basis of previous studies that evaluated inhibition of other viral infections (15,16), as well as studies of its role as an effective anticancer cardiac glycoside (19). Because oleandrin was dissolved in DMSO, negative controls containing equal amounts of DMSO in the absence of oleandrin were also tested. First, a lactate dehydrogenase release (LDR) assay for cytotoxicity was performed to establish the suitability of the intended experimental system. Overall, oleandrin was well tolerated by the highly SARS-C-V-2-susceptible Vero CCL81 cell line (20)(Fig. 1). At 24 hours post-treatment, no significant differences were observed between oleandrin and the DMSO controls (Fig. 1A,B). By 48 hours post-treatment, significant differences were observed (Fig. 1C,D). However, the absolute differences were relatively modest compared to the maximum cytotoxicity of Triton-X100, which was included as an assay control. The highest cytotoxicity of oleandrin was observed at the 0.1μg/ml concentration, with an increase of 1.86% cytotoxicity in the DMSO control to 6.88% in the oleandrin-treated wells for a drug-dependent difference of 5.02%. All other oleandrin concentrations caused less than 2% overall cytotoxicity.

**Figure 1.**
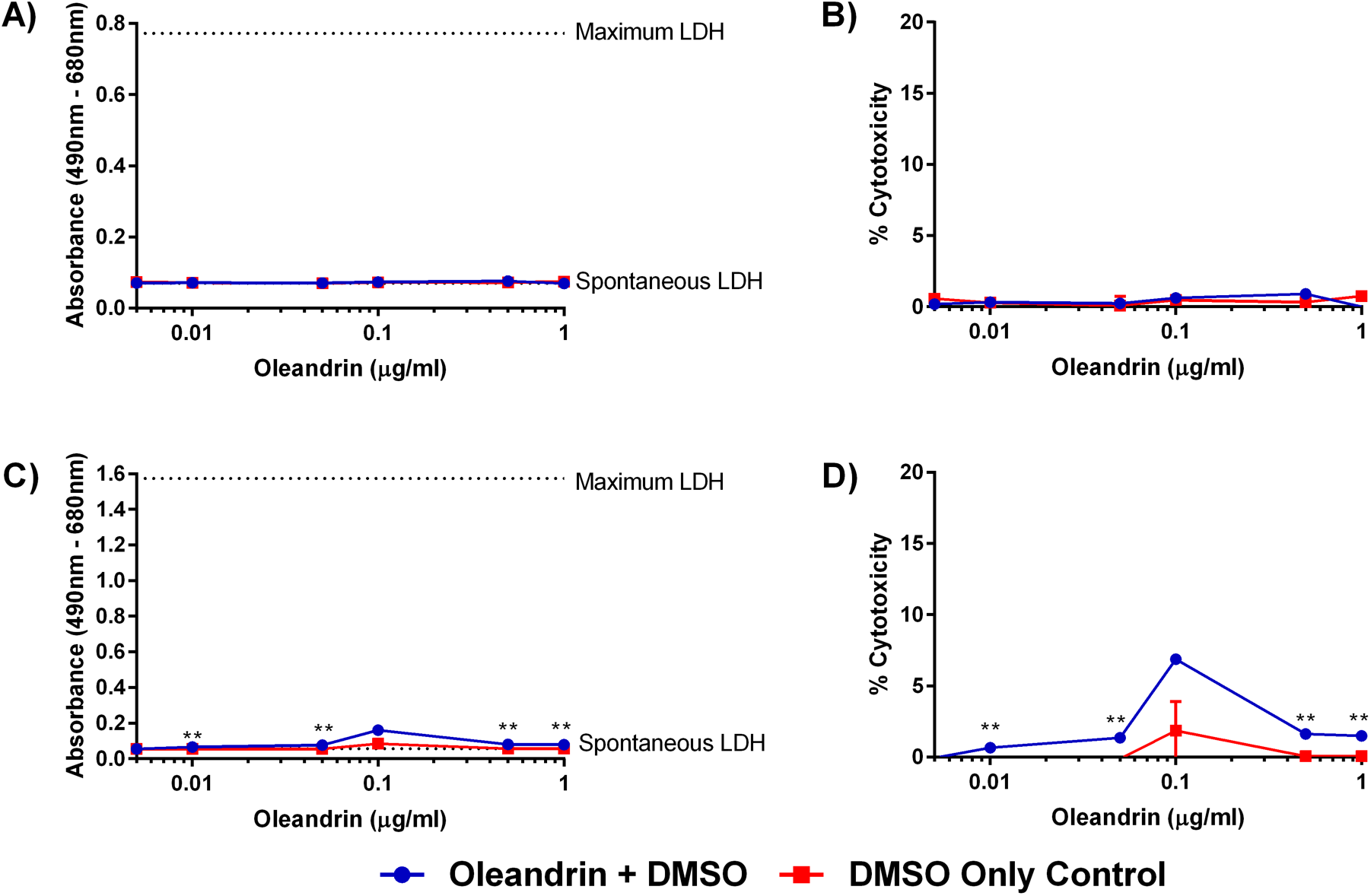
Oleandrin is well tolerated by Vero CCL81 cells. Release of LDH was measured as a proxy for cytotoxicity potentially induced by varying concentrations of oleandrin and DMSO-matched controls. LDH levels released in the supernatant were measured at (A and B) 24 hours and (C and D) 48 hours after the addition of oleandrin. Raw absorbance values (A and C) were normalized to a percent cytotoxicity (B and D), with the spontaneous release of LDH in untreated cells set as 0 and the maximum release of LDH in Triton-X100-treated cells set as 100. Symbols (circles, squares) represent the mean, error bars represent standard deviations. Oleandrin-treated cells and their DMSO-matched controls were compared via T-tests with Holms correction for multiple comparisons to test for any drug-associated cytotoxicity. ** = p<0.01.

### Prophylactic Efficacy of Oleandrin

To assess the prophylactic potential of oleandrin in the context of SARS-CoV-2 infection, Vero CCL81 cells, shown previously to be highly susceptible (20), were pre-treated with oleandrin or DMSO-carrier controls for 30 minutes prior to infection. SARS-CoV-2, strain USA_WA1/2020, a 4^th^ passage isolate from a patient treated in Washington State, U.S. (20), was used to infect the cells, and post-infection culture medium was also supplemented with either oleandrin in DMSO, or DMSO alone. Culture supernatant samples were collected at 24 and 48 hours post-infection, at which point the experiments were terminated due to the onset of substantial viral-associated cytopathic effects (CPE) in the DMSO controls.

In the absence of oleandrin, SARS-CoV-2 reached approximately 6 log_10_ plaque-forming units (pfu)/ml titers by the 24-hour timepoint, and maintained that titer at the later timepoint (Fig. 2 A,B). The DMSO controls all reached similar titers, with no dose-dependent trend observed (Supplemental Figure 1). The four highest concentrations of oleandrin (1.0 μg/ml – 0.05 μg/ml) significantly reduced the production of infectious SARS-CoV-2. The 1.0 μg/ml and 0.5 μg/ml oleandrin concentrations both reduced SARS-CoV-2 titers by more than 4 log_10_ pfu/ml, which consistently remained either at or below the limit of detection for the assay. This corresponded to an approximately 20,000-fold reduction in infectious SARS-CoV-2 titer for the 1.0 μg/ml and 0.5 μg/ml oleandrin concentrations (Fig. 2C). The 0.1 μg/ml oleandrin dose resulted in a greater than 3,000-fold reduction, and the 0.05 μg/ml concentration resulted in an 800-fold reduction. The 0.01 μg/ml and 0.005 μg/ml concentrations had no significant effect on SARS-CoV-2 production.

**Figure 2.**
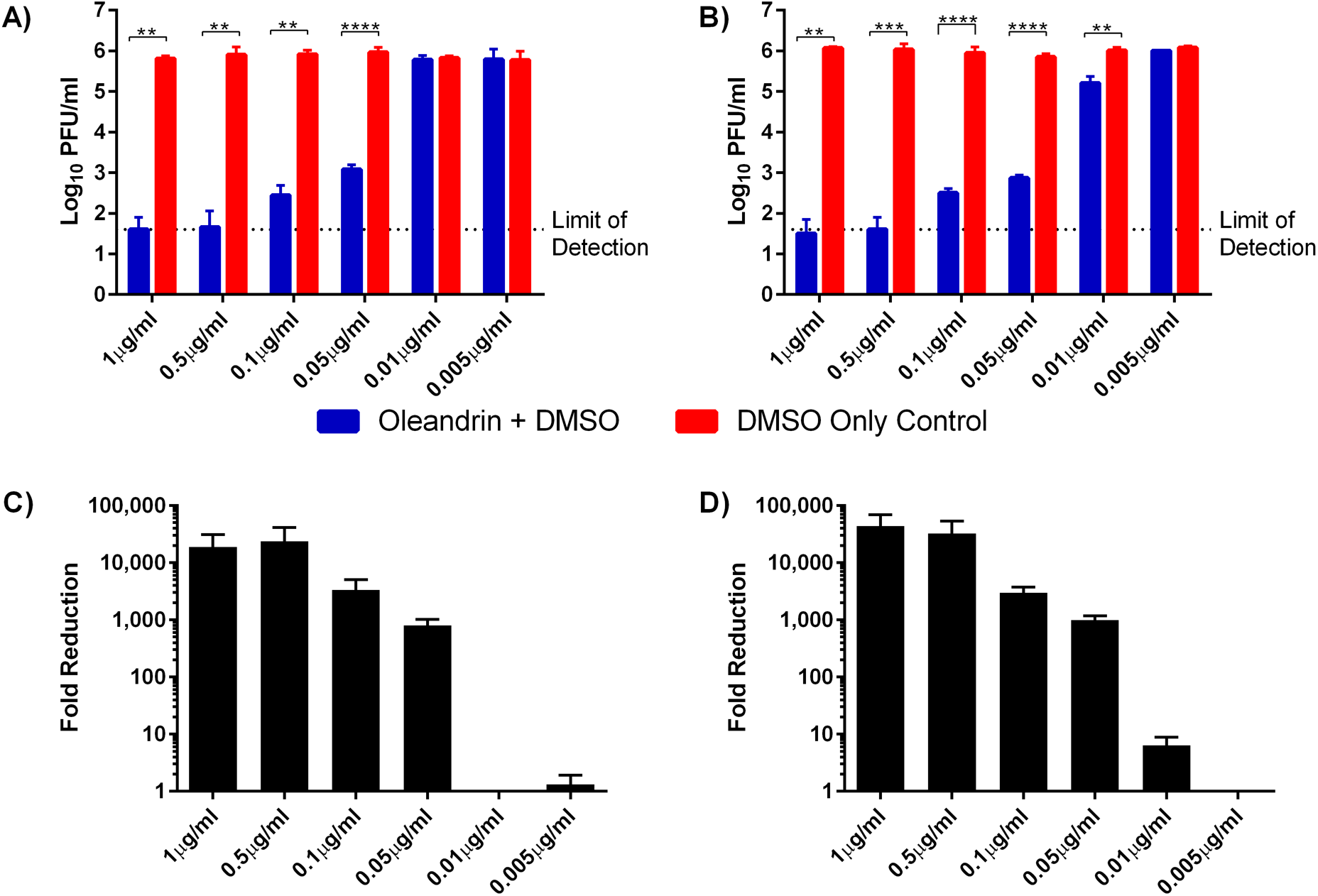
Oleandrin prophylactically reduces SARS-CoV-2 infectious titer when added prior to infection. Various concentrations of oleandrin or DMSO-matched controls were added to Vero CCL81 cells 30 minutes prior to infection (MOI = 0.01), and were maintained after infection as well. Supernatant was collected (A and C) 24 hours and (B and D) 48 hours after infection. Infectious SARS-CoV-2 titers were quantified via plaque assay (A and B), and the fold reduction in oleandrin-treated samples vs. their corresponding DMSO-matched controls were calculated (C and D). Data shown from a single representative experiment conducted in triplicate. Bar heights represent the mean, error bars represent the standard deviation. Oleandrin treated samples and DMSO-matched controls were compared via t tests, with a Holms correction for multiple comparisons. ** = p<0.01, *** = p<0.001, **** = p<0.0001.

At the 48-hour timepoint, the DMSO controls maintained approximately the same titer observed earlier, while the 1.0 μg/ml – 0.05 μg/ml oleandrin concentrations maintained their significant reductions in SARS-CoV-2 titers. Interestingly, the 0.01 μg/ml dose, which had no significant effect compared to its DMSO control at 24 hours post-infection, did result in a statistically significant reduction in titer at 48 hours post-infection (p<0.01). This trend of a greater drug-induced reduction observed at 48 hours as compared to 24 hours, was consistent at the higher doses of oleandrin as well (Fig. 2C, D). At 1.0 μg/ml, the effect increased to from 18,822-fold reduction at 24 hours to 44,000-fold reduction at 48 hours, and at 0.5 μg/ml the effect increased from 23,704-fold to 32,278-fold. The reductions observed for 0.1 μg/ml and 0.05 μg/ml remained similar at 24 and 48 hours post-infection. The 0.01 μg/ml concentration of oleandrin decreased SARS-CoV-2 titers by 6.3-fold. The increased prophylactic efficacy of oleandrin over time (24 vs. 48 hours) was reflected in its EC_50_ values, calculated at 11.98ng/ml at 24 hours post-infection and 7.07ng/ml at 48 hours post-infection.

The results of this prophylactic study were confirmed in a second, independent experiment (Supplementary Figure 2).

### Genomic Equivalents and Genome:PFU Ratios

Cardiac glycosides have been shown to inhibit the formation of infectious HIV particles through disruption of the virus envelope (15). To determine whether the inhibition of SARS-CoV-2 was at the level of total or infectious particle production, RNA was extracted from the cell culture supernatants of the prophylactic study and genomic equivalents were quantified via RT-qPCR (Fig. 3). The prophylactic effect of oleandrin, initially observed via infectious assay, was confirmed at the level of genome equivalents. At 24 hours-post infection, oleandrin significantly decreased SARS-CoV-2 genomes in the supernatant at the four highest doses (Fig. 3A). By 48 hours post-infection, all but the lowest dose resulted in significant reductions in SARS-CoV-2 genomes (Figure 3B), confirming the results observed from the infectious assay in which the 0.01 μg/ml concentration gained efficacy as the infection proceeded.

**Figure 3.**
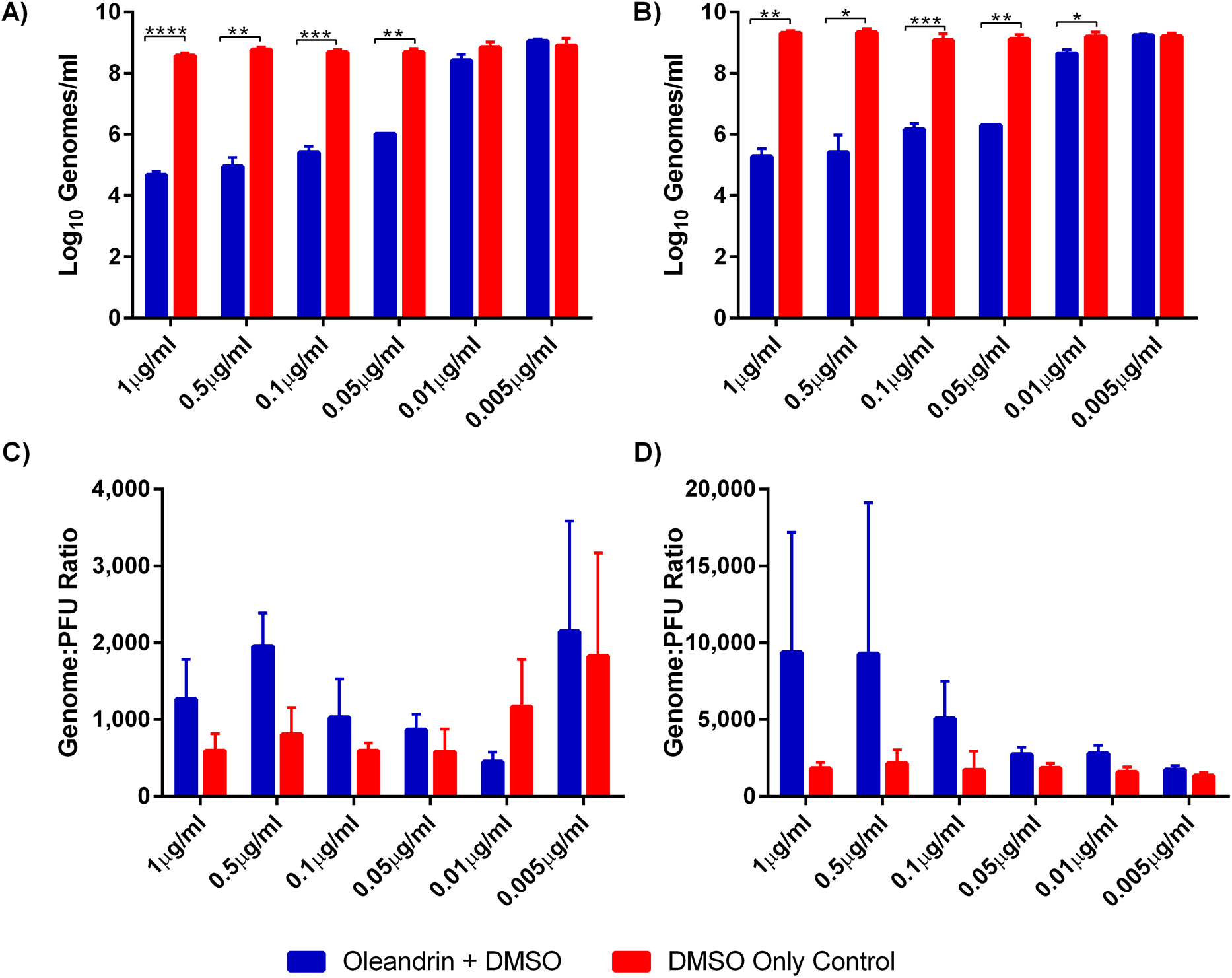
Oleandrin prophylactically reduces SARS-CoV-2 genomes when added prior to infection. Various concentrations of oleandrin or DMSO-matched controls were added to Vero CCL81 cells 30 minutes prior to infection (MOI = 0.01), and were maintained after infection as well. Supernatant was collected (A and C) 24 hours and (B and D) 48 hours after infection. SARS-CoV-2 genomes were quantified via qRT-PCR (A and B), and the genome to PFU ratios were calculated using the previous obtained infectious titers (C and D). Data shown from a single representative experiment conducted in triplicate. Bar heights represent the mean, error bars represent the standard deviation. Oleandrin treated samples and DMSO-matched controls were compared via T tests, with a Holms correction for multiple comparisons. * = p<0.05, ** = p<0.01, *** = p<0.001, **** = p<0.0001.

Genome:infectious particle ratios were also calculated (Figs. 3C,D). While there were no significant differences between the oleandrin-treated samples and their DMSO controls, there were several interesting trends. First, at both 24 and 48 hours post-infection the higher, more efficacious doses of oleandrin had higher genome:pfu ratios than their DMSO controls (Figs. 3 C,D). This effect was more pronounced at 48 hours than at 24 hours post-infection, suggesting that oleandrin may have reduced infectivity of viral progeny.

### Therapeutic Efficacy of Oleandrin

In addition to the prophylactic study, the effect of oleandrin when added after SARS-CoV-2 infection was measured to simulate a therapeutic application. The 1.0 μg/ml – 0.05 μ/ml concentrations, which consistently demonstrated a prophylactic effect in the above experiments, were added at either 12 or 24 hours post-infection. DMSO-matched controls were again included. Samples were harvested from the 12-hour treatment at both 24 and 48 hours post-infection (Figs. 4A-D). At 24 hours-post infection, oleandrin reduced SARS-CoV-2 titers by approximately 10-fold (Fig. 4B), despite a short contact time of only 12 hours. Importantly, by 48 hours post-infection, the therapeutic effect became much more pronounced both in terms of the titer reduction and absolute titers. The highest dose resulted in a greater than 1,000-fold reduction in infectious SARS-CoV-2 titer, with the 0.5 μg/ml and 0.1 μg/ml doses causing greater than 100-fold reductions, and the 0.05 μg/ml dose resulting in a 78-fold reduction (Fig. 4D).

**Figure 4.**
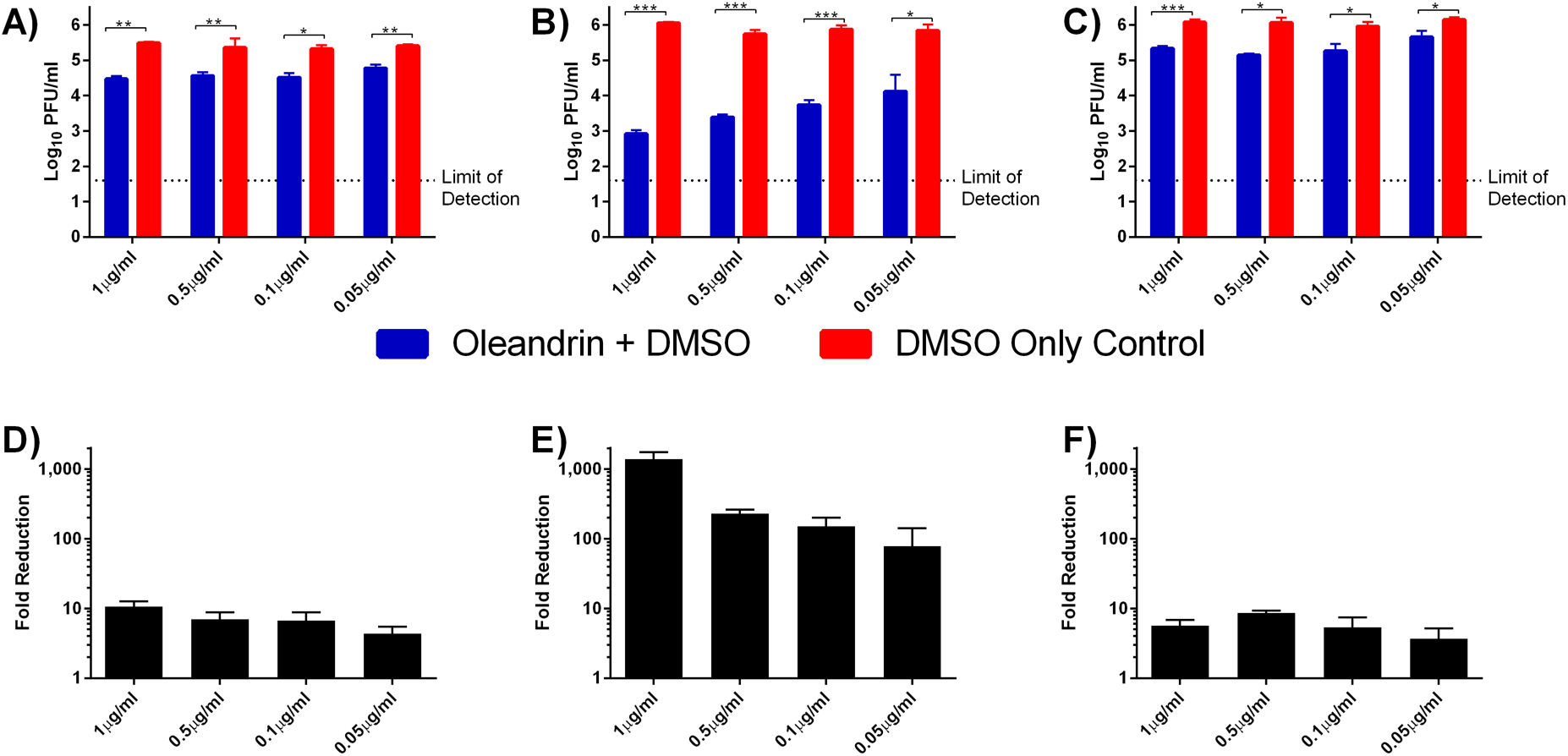
Oleandrin reduces SARS-CoV-2 infectious titer when added after infection. Various concentrations of oleandrin or DMSO-matched controls were added to Vero CCL81 cells either 12 hours (A, B, D, and E) or 24 hours (C and F) after infection (MOI = 0.01). Supernatant was collected (A and D) 24 hours after infection from the samples previously drugged at 12 hours post-infection. Supernatant was also collected 48 hours after infection from both the samples previously drugged at 12 hours post-infection (B and E) and at 24 hours post-infection (C and F). Infectious SARS-CoV-2 titers were quantified via plaque assay (A, B, and C), and the fold reduction in oleandrin-treated samples vs. their corresponding DMSO-matched controls were calculated (D, E, and F). Data shown from a single representative experiment conducted in triplicate. Bar heights represent the mean, error bars represent the standard deviation. Oleandrin treated samples and DMSO-matched controls were compared via t tests, with a Holms correction for multiple comparisons. * = p<0.05, ** = p<0.01, *** = p<0.001.

Oleandrin maintained its therapeutic effect even when added at 24 hours post-infection (Fig. 4E,F), a timepoint at which extensive SARS-CoV-2 replication has already occurred. The 24-hour treatment resulted in a 5-10-fold reduction in titers measured at 48 hours post-infection, similar to the impact of 12-hour treatment measured at 24 hours post-infection. Unfortunately, it was not possible to test time points beyond 48 hours to determine the extended therapeutic efficacy due to the viral-induced CPE in the DMSO-only control wells.

## Discussion

The emergency COVID-19 pandemic situation calls for rapid screening of potential therapeutic or prophylactic drugs already shown in prior clinical trials to be safely tolerated. Here, we tested the prophylactic and therapeutic *in vitro* efficacy of oleandrin, a cardiac glycoside extracted from the *Nerium oleander* plant. As in our previous studies with other viruses, we found that oleandrin exhibits potent antiviral activity against SARS-CoV-2. Prophylactic treatment of Vero cells with as little as 0.05 μg/ml resulted in a significant, 800-fold reduction in virus production, and the 0.1 μg/ml oleandrin dose resulted in a greater than 3,000-fold reduction in infectious virus as well as a similar reduction in viral RNA. The EC_50_ values were 11.98ng/ml when virus output was measured at 24 hours post-infection, and 7.07ng/ml measured at 48 hours post-infection. Therapeutic (post-infection) treatment up to 24 hours after infection of Vero cells also reduced viral titers, with the 0.5 μg/ml and 0.1 μg/ml doses causing greater than 100-fold reductions measured at 48 hours, and the 0.05 μg/ml dose resulting in a 78-fold reduction.

The pharmacology of oleandrin has been a subject of expanded examination during the past 20 years (21). Its most recognized activity is inhibition of Na,K-ATPase through blockade of ATP binding sites. Other membrane-positioned enzymes requiring ATP include ACE-2 which has now been shown to be an entry receptor for SARS-CoV-2 (22). The potential role of oleandrin as an active principle ingredient in the botanical drug (PBI-05204) designed to treat malignant disease has been reported (12–14, 19). The Phase I trial of PBI-05204, for example, included measurement of Cmax and t_1/2_ calculations for oleandrin after oral administration to cancer patients. The delayed oral administration of this candidate drug achieved plasma concentrations of 2-4 ng oleandrin/ml, without significant adverse effects, at a dose determined to be appropriate for Phase II study. The measured half-life was variable at 8-13 hr, indicating that multiple daily administrations may be optimal for maximum benefit. Our unpublished studies have shown that sublingual administration of oleandrin as an extract of *Nerium oleander* offers facile, safe and rapid plasma concentrations of oleandrin that surpass those achieved after oral administration. This is not surprising because oral administration of oleandrin results in cleavage of the molecule in liver tissue to the much less active metabolite oleandrigenin.

Prior preclinical research demonstrated that oleandrin, in contrast to other cardiac glycosides such as digoxin, readily crosses the blood-brain barrier, where it induces beneficial effects such as the induction of brain derived neurotrophic factor (BDNF) and induction of nuclear factor erythroid 2-related factor 2 (Nrf-2), providing antioxidant activities (23). Given the poorly understood neurological manifestations presenting in some COVID-19 patients (24, 25), oleandrin may therefore be of some benefit in preventing virus-associated neurological disease. Of course, further studies are needed including in animal models that reproduce neurological signs.

Oleandrin has also been shown to produce a strong anti-inflammatory response, which may be of benefit in preventing hyper-inflammatory responses to infection with SARS-CoV-2 (26). We show here that low concentrations of oleandrin produce strong antiviral activity against SARS-CoV-2. Thus, considering the beneficial anti-inflammatory activity of *Nerium oleander* extracts combined with the suggestion that oleandrin might protect against neurological deficits, Phoenix Biotechnology, Inc. (San Antonio, TX) has developed an extract for sublingual administration that it will quickly explore for efficacy in appropriate *in vivo* studies.

Additional studies are needed to define the mechanism of oleandrin’s antiviral activity. Several steps in viral replication could be affected including entry/fusion, lipid metabolism related to budding, formation of replication complexes, secretory pathway function and affects on SARS-CoV-2 protein processing, etc. Also, in vivo studies with animal models such as transgenic mice (27), hamsters (28), ferrets (29) or nonhuman primates (30) should be conducted, and oleandrin should be tested for antiviral activity against SARS-CoV and MERS-CoV.

In summary, we demonstrated that oleandrin has potent anti-SARS-CoV-2 activity *in vitro*, both when administered before or after infection. Care must be taken when inferring potential therapeutic benefits from *in vitro* antiviral effects. Nevertheless, our results indicate the *in vivo* studies with oleandrin should proceed rapidly to address the COVID-19 pandemic.

## Methods

### Cells, Viruses, and Oleandrin

Vero CCL81 cells were used for the prophylactic and therapeutic assays (ATCC, Manassas, VA). Plaque assays were performed in Vero E6 cells, kindly provided by Vineet Menachery (UTMB, Galveston, TX). The cells were maintained in a 37°C incubator with 5% CO_2_. Cells were propagated utilizing a Dulbecco’s Modified Eagle Medium (Gibco, Grand Island, NY) supplemented with 5% fetal bovine serum (FBS) (Atlanta Biologicals, Lawrenceville, GA) and 1% penicillium/streptomycin (Gibco, Grand Island, NY). Maintenance media reduced the FBS to 2%, but was otherwise identical.

SARS-CoV-2, strain USA_WA1/2020 (Genbank accession MT020880), was provided by the World Reference Center for Emerging Viruses and Arboviruses (20). All studies utilized a NextGen sequenced Vero passage 4 stock of SARS-CoV-2.

Oleandrin (PhytoLab, Vestenbergsgreuth, Germany) was dissolved at a concentration of 1mg/ml in DMSO (Invitrogen, Eugene, OR).

### LDH Toxicity Assay

96-well plates were seeded with 2×10^4^ Vero CCL81 cells in 100ul growth media and returned to a 37C/5%CO_2_ incubator overnight. The following day, the growth media was removed and replaced with 100ul maintenance media containing either 1-0.005 ug/ml oleandrin in 0.1-0.0005% DMSO, 0.1-0.0005% DMSO without drug, or untreated media to serve as controls for the maximum and spontaneous release of LDH. All treatments were added to triplicate wells, and the plate was returned to the 37C/5%CO_2_ incubator. At either 24 or 48 hours post-treatment, the plate was removed from the incubator and the LDH released into the supernatant was assessed CyQUANT LDH toxicity assay (Invitrogen, Eugene, OR) according to the manufacturer’s directions. Absorbance was measured at 490nm and 680nm on a Synergy HT plate reader (BioTek, Winooski, VT) with Gen5 data analysis software, version 2.05 (BioTek, Winooski, VT).

### Prophylactic and Therapeutic Studies

In the prophylactic studies, growth media was removed from confluent monolayers of approximately 10^6^ Vero CCL81 cells in 6-well plates. The growth media was replaced with 200μl of maintenance media containing either 1.0 μg/ml, 0.5μg/ml, 0.1 μg/ml, 0.05 μg/ml, 0.01 μg/ml, or 0.005 μg/ml oleandrin, or matched DMSO-only controls. Treated plates were returned to the 37°C/5%CO_2_ incubator for thirty minutes. After pre-treatment, 1 × 10^4^ SARS-CoV-2 plaque forming units (PFU) in a volume of 200μl maintenance media was added to all wells, yielding a multiplicity of infection of 0.01. The infection was allowed to proceed for one hour at 37°C and 5% CO_2_. After infection, cells were gently washed 3 times with DPBS (Sigma, St Louis, MO). Finally, 4ml of maintenance media containing either oleandrin in DMSO or DMSO-only was added to each well according to the pre-treatment concentrations. Samples were taken 24- and 48-hours post-infection. Each dose and control pair were done in triplicate, and the entire study was repeated, again in triplicate.

In the therapeutic studies, the pre-treatment step was eliminated. Vero CCL81 cells were seeded, infected, washed, and returned to the incubator as before. At either 12- or 24-hours post-infection, oleandrin in DMSO or matched DMSO-only controls was spiked into the appropriate wells. The two lowest doses (0.01 μg/ml and 0.005 μg/ml) were not included. Samples from the 12 hour post-infection treatment were taken 24 and 48 hours post-infection. Samples from the 24 hour post-infection treatment were taken at 48 hours post-infection. Each dose and control pair were done in triplicate.

### Plaque Assay

Vero E6 cells were seeded in 6-well plates (Thermo Scientific, Waltham, MA) to a confluency of approximately 90%. Samples were serially 10-fold diluted in DPBS. Diluted samples were plated onto the cells with a volume of 0.25ml and left to incubate at 37°C with 5% CO_2_ for one hour. After incubation, samples were overlaid with a 1:1 mixture of one part 2X MEM (Gibco, Grand Island, NY) supplemented with 8% FBS and 2% pen/strep, and one part 1.6% LE agarose from (Promega, Madison, WI). The plates were incubated for two days at 37°C at 5% CO_2_. Plaques were visualized with neutral red stain (Sigma, St Louis, MO) and a light box.

### Genome copy measurement

To quantify genome copies for the samples, 200μl of sample was extracted with a 5:1 volume ratio of TRIzol LS (Ambion, Carlsbad, CA), utilizing standard manufacturers protocols and resuspending in 11μl water. Extracted RNA were tested for SARS-CoV-2 by qRT-PCR following a previously published assay (26). Briefly, the N gene was amplified using the following primers and probe: forward primer [5’-TAATCAGACAAGGAACTGATTA-3’]; reverse primer [5’-CGAAGGTGTGACTTCCATG-3’]; and probe [5’-FAM-GCAAATTGTGCAATTTGCGG-TAMRA-3’]. A 20μl reaction mixture was prepared using the iTaq Universal probes One-Step kit (BioRad, Hercules, CA), according to manufacturer instructions: A reaction mix (2x: 10μL), iScript reverse transcriptase (0.5 μL), primers (10μM: 1.0 μL), probe (10μM: 0.5 μL), extracted RNA (4μL) and water (3 μL). The RT-qPCR reactions were conducted using the thermocycler StepOnePlus™ Real-Time PCR Systems (Applied Biosystems). Reactions were incubated at 50°C for 5min and 95°C for 20sec followed by 40 cycles at 95°C for 5sec and 60°C for 30sec. The positive control RNA sequence (nucleotides 26,044 - 29,883 of COVID-2019 genome) was provided by Dr. Pei-Yong Shi and used to estimate the RNA copy numbers of N gene in the samples under evaluation.

### Statistical Analysis

Matched Oleandrin-treated samples and DMSO-only controls had their LDH absorbance, percent cytotoxicity, Log_10_-transformed infectious and genomic equivalent titers, and raw genome to PFU ratios analyzed via t-tests (one per concentration) with Holm’s correction for multiple comparisons. Values below the limit of detection were assumed to equal one-half of that value for graphing and statistical purposes. All graphs were generated in Prism version 7.03 (GraphPad, San Diego, CA). All statistical analysis was performed in R version 3.6.2 (27).

## Acknowledgments

This research was supported by NIH grant R24 AI120942 to SCW. We thank Natalie Thornburg (Centers for Disease Control and Prevention, Atlanta, GA, USA) for providing the SARS-CoV-2 USA_WA1/2020 isolate.

## Competing interests

RAN is Chief Science Officer of Phoenix Biotechnology Inc.

KJS is a paid consultant with Phoenix Biotechnology Inc.

---

**Supplementary Figure 1.**
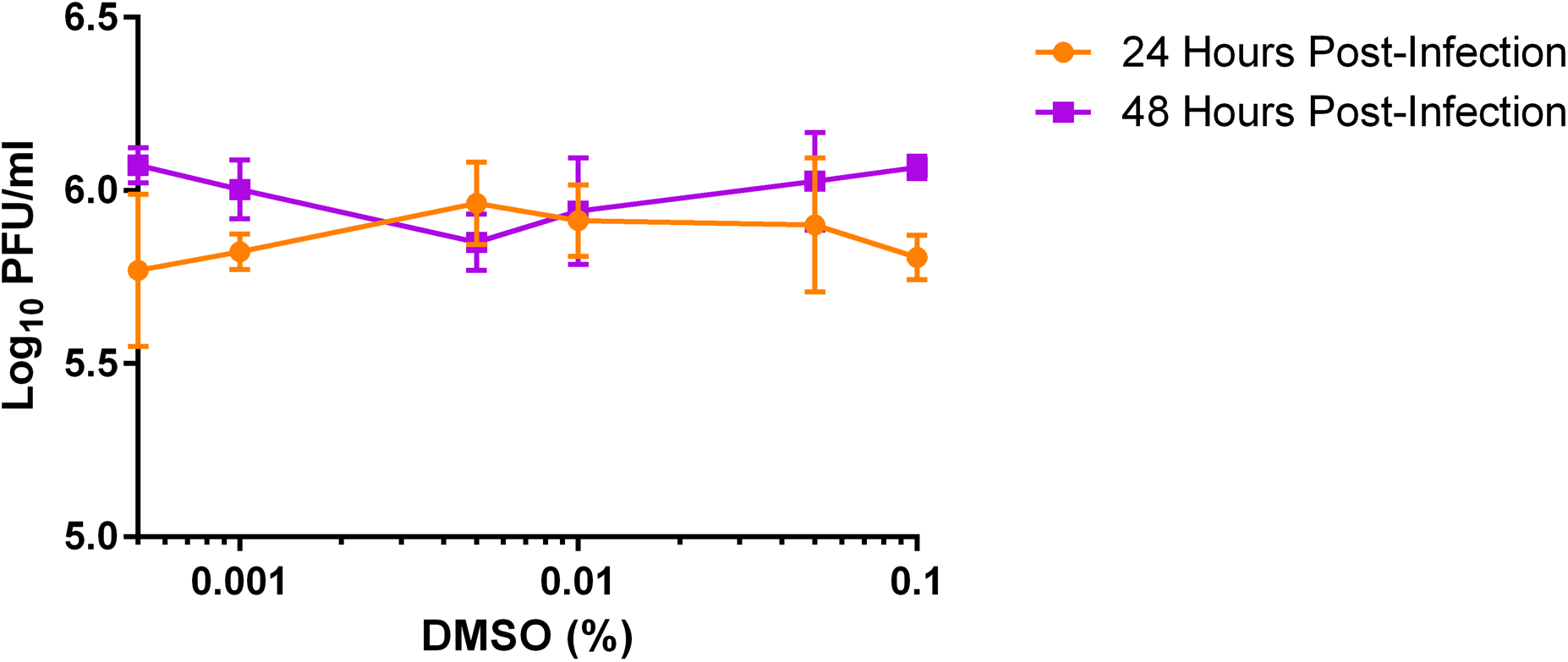
DMSO vehicle does not affect the replication of SARS-CoV-2. To control for the possible impact of DMSO on SARS-CoV-2 replication and infectious titer (MOI = 0.01), DMSO-matched controls were included alongside oleandrin. The concentrations of DMSO used in this study (0.1% - 0.0005%) had no apparent impact on SARS-CoV-2 titer at either 24 or 48 hours post-infection. No dose-dependent trend was apparent, and ANOVA revealed no statistically significant variation (p = 0.53449 at 24 hours and p = 0.12697 at 48 hours). Symbols (circles, squares) represent the mean, error bars represent the standard deviation. Data shown from a single representative experiment conducted in triplicate.

**Supplementary Figure 2.**
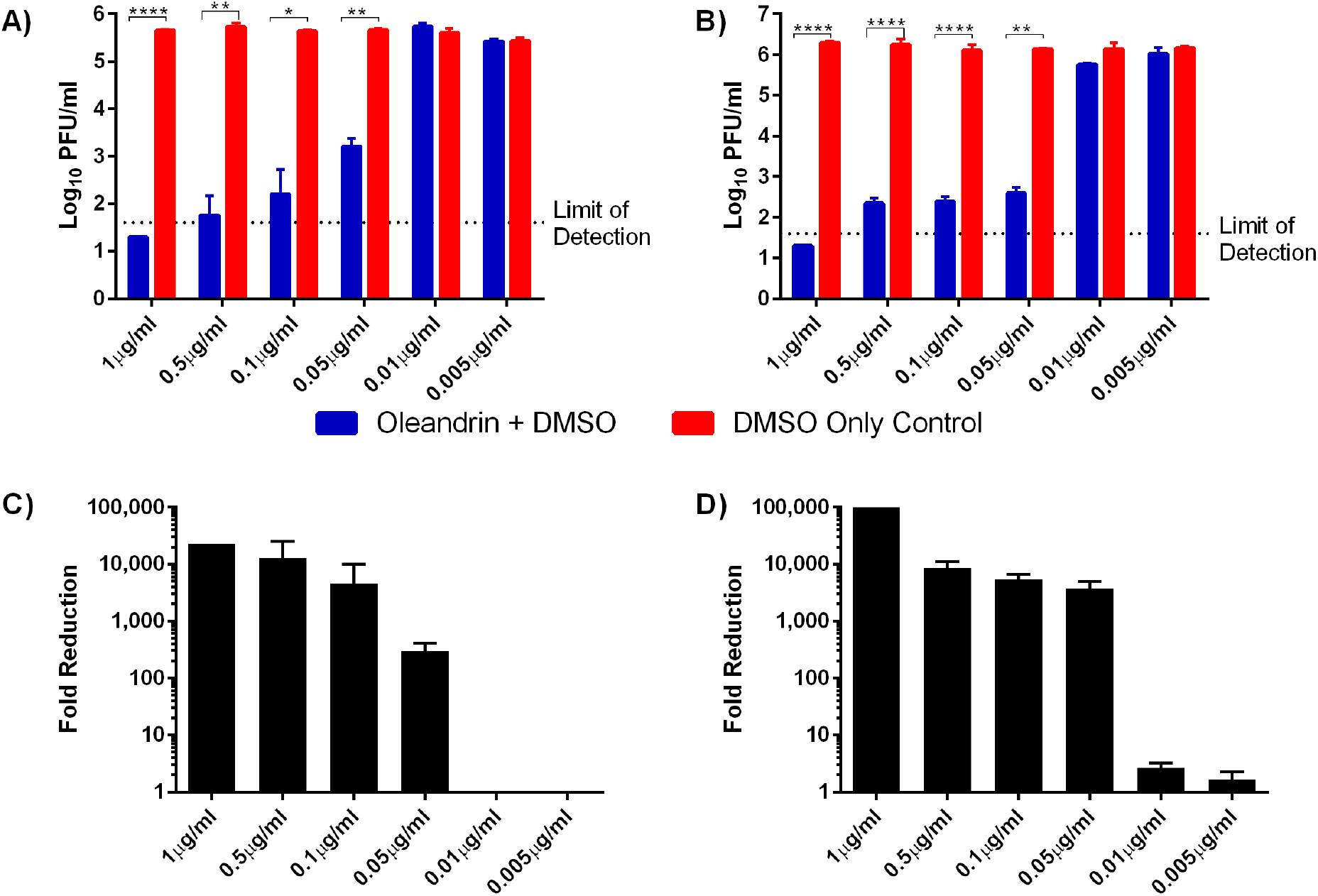
The prophylactic efficacy of oleandrin is highly repeatable. A separate, independent study was undertaken to confirm the prophylactic efficacy of oleandrin. As before, various concentrations of oleandrin or DMSO-matched controls were added to Vero CCL81 cells 30 minutes prior to infection (MOI = 0.01), and were maintained after infection as well. Supernatant was collected (A and C) 24 hours and (B and D) 48 hours after infection. Infectious SARS-CoV-2 titers were quantified via plaque assay (A and B), and the fold reduction in oleandrin-treated samples vs. their corresponding DMSO-matched controls were calculated (C and D). Data shown from a single representative experiment conducted in triplicate. Bar heights represent the mean, error bars represent the standard deviation. Oleandrin treated samples and DMSO-matched controls were compared via t tests, with a Holms correction for multiple comparisons. * = p<0.05, ** = p<0.01, **** = p<0.0001.

